# Nucleoprotein phase-separation affinities revealed via atomistic simulations of short peptide and RNA fragments

**DOI:** 10.1101/2024.09.24.614800

**Authors:** Vysakh Ramachandran, William Brown, Davit A Potoyan

## Abstract

Liquid-liquid phase separation of proteins and nucleic acids into condensate phases is a versatile mechanism for ensuring compartmentalization of cellular biochemistry. RNA molecules play critical roles in these condensates, particularly in transcriptional regulation and stress responses, exhibiting a wide range of thermodynamic and dynamic behaviors. However, deciphering the molecular grammar that governs the stability and dynamics of protein-RNA condensates remains challenging due to the multicomponent and heterogeneous nature of these biomolecular mixtures. In this study, we employ atomistic simulations of twenty distinct mixtures containing minimal RNA and peptide fragments to dissect the phase-separating affinities of all twenty amino acids in the presence of RNA. Our findings elucidate chemically specific interactions, hydration profiles, and ionic effects that synergistically promote or suppress protein-RNA phase separation. We map a ternary phase diagram of interactions, identifying four distinct groups of residues that promote, maintain, suppress, or disrupt protein-RNA clusters.

---

RNA molecules in cells are seldom found in isolation; they typically associate with proteins to form ribonucleoprotein complexes (RNPs). These complexes subsequently localize within various membrane-less organelles or compartments, driven by the mechanism of liquid-liquid phase separation (LLPS). Notable examples of RNA-rich membraneless organelles include nucleoli, nuclear speckles, and stress granules, which form in response to cellular stress and play a critical role in mRNA metabolism and storage [1–3]. RNAs are often seen as the drivers of phase separation thanks to their ability to form multivalent interactions with other flexible RNAs or proteins containing low-complexity domains (LCDs). In vitro experiments with LCD sequences have shown great tunability of physicochemical and material properties of RNA-protein condensates towards sequence variations in LCDs [4–6]. For instance, single residue substitutions in repeat peptide sequences have been shown to condensate droplet fusion times dramatically later, selective partitioning of small molecules, viscoelastic dynamics, and maturation time scales [2,3,7]. While recent experimental and computational studies have primarily explored the roles of positively charged aromatic residues and the importance of multivalent cation-π and π-π interactions[8–11], a comprehensive understanding of how all twenty amino acid residues interact with RNA to influence condensate properties remains incomplete.

The thermodynamic impact of LCD sequence variations is commonly rationalized by employing the sticker-spacer framework inspired by the physics of associative polymers [12,13]. Most LCDs are enriched with residues classified as stickers interspersed by spacer regions, which allow the LCD to maximize interactions with RNAs. Specific sticker residues, such as tyrosine [14], tryptophan [15], arginine [6,10,14,16], serine [14,17], glutamine [14,18], and asparagine [17,18], have been identified as critical determinants in promoting RNA binding protein (RBP) condensation with RNA, based on their specific interactions with RNA.

However, both experiments[6,16,19] and simulations[4,11,20] suggest that a simple classification of sticker versus spacer interactions may only partially explain the complex hierarchy of forces governing phase-separation affinities of LCDs toward RNA. A more nuanced approach may be necessary to rationalize these interactions and their influence on phase behavior, including predicting material properties and heterogeneous multiphasic morphologies[2,21]. Furthermore, the role of molecules of RNA is shown to undergo homotypic phase separation[22] in the presence of divalent ions, suggesting that complex ion and solvent-mediated RNA-RNA associations can be ignored. Understanding the molecular sequence grammar that governs protein-RNA LLPS can pave the way for tuning the biophysical characteristics of condensates. This tuning could potentially serve as a therapeutic intervention technique to prevent the irreversible liquid-to-solid transitions [23] that accompany various neurodegenerative conditions, such as amyotrophic lateral sclerosis (ALS), frontotemporal dementia (FTD), Alzheimer’s, and Parkinson’s [24–27].

In this work, we employ atomistic simulations of twenty different mixtures containing minimal tri-uracyl and tri-peptide fragments to disentangle phase-separating affinities of RNA towards all twenty amino acids.

We have chosen the minimal sticker motifs of GXG tripeptides for peptide fragments where the central residue X is varied. For RNA fragments, we have chosen tri-uracil [U]_3_. Each mixture contains 1:1 mass ratio of distinct GXG peptide fragment with RNA fragment which ensures ∼100 mg/ml biomolecular density and approximately 3:1 stochiometric ratio of protein to RNA. Previous work on peptide phase-separation[28,29] has shown that 50-150 mg/ml is optimal density for observing phase-separation and formation of clusters (See Methods and SI). We have simulated each mixture for over two microseconds, which exceeds cluster formation and reorganization time by order of magnitude. Cluster formation could be seen as early as ∼200 ns. Globally, the simulations show the rapid formation of GXG-RNA complexes, which vary in size and density depending on the nature of the peptide in the mixtures. We quantified the densities and distributions of cluster sizes and a fraction of peptides in each cluster, which reflects the propensity of each central residue to phase separate in the presence of RNA (Figure 1B-D).

**Figure 1.**
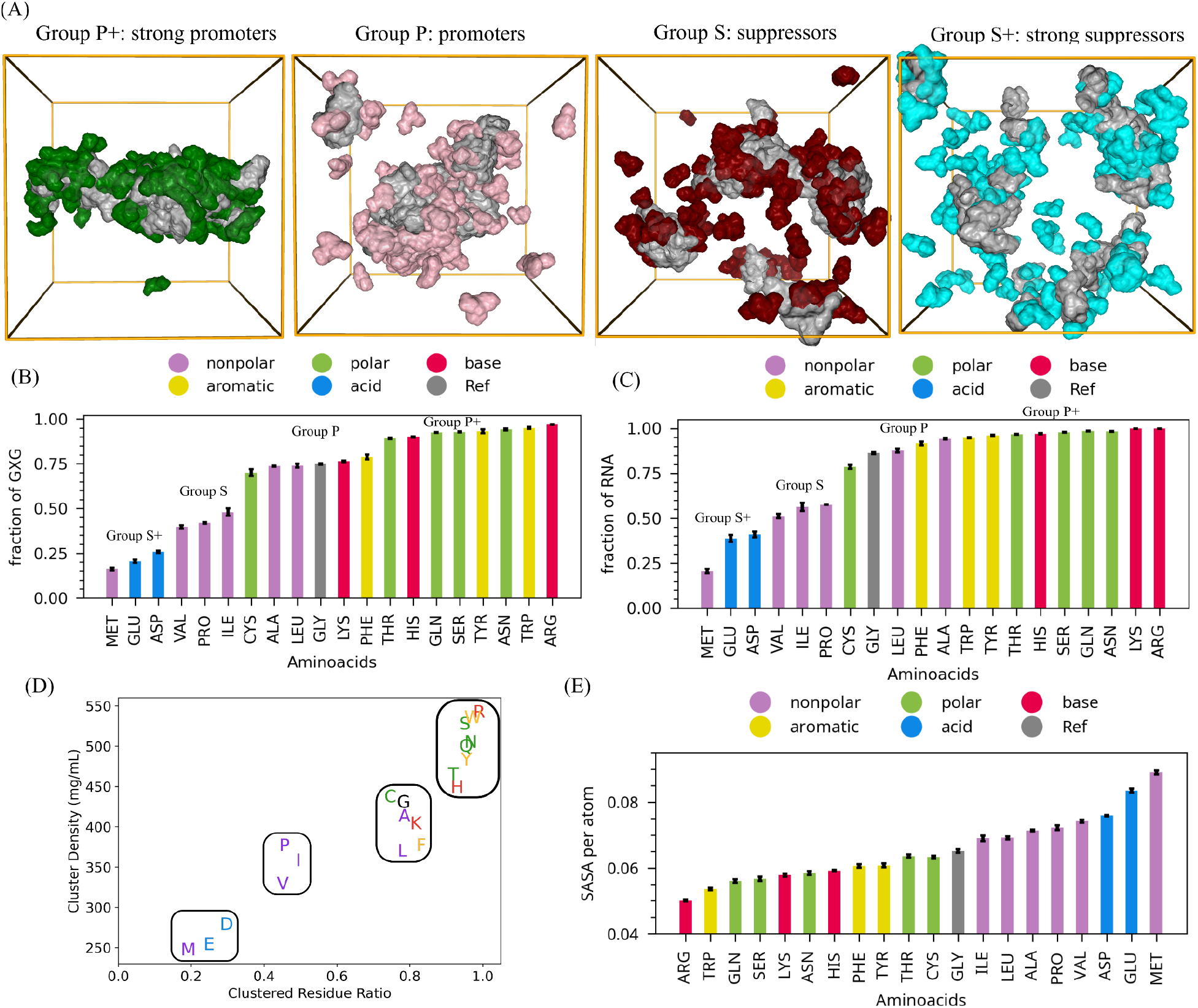
Condensation propensity of central X residues in GXG peptides with Uracil bases of RNA. (A) Representative snapshots of simulations for four groups based on their condensation propensity: Group P+: strong promoter (ARG), Group P: promoter (PHE), Group S: suppressor (PRO), and Group S+: strong suppressor (MET). Cluster size is quantified by (B) the fraction of GXG peptides and (C) the fraction of PolyU fragments inside the cluster. (D) Cluster density vs. cluster size for all 20 residues. (E) SASA per atom in the cluster for each system.

The four distinct groups can be clearly identified when plotting the density of the cluster vs the fraction of peptides in the cluster (Figure 1A-D). We defined the four groups as follows; Group P+: strong promoters, including ARG, TRP, ASN, TYR, SER, GLN, HIS, and THR. Group P: promoters, including CYS, GLY, LEU, LYS, PHE, and ALA; Group S: suppressors, including VAL, PRO, and ILE; and Group S+: strong suppressors, including MET, ASP, and GLU. The fraction of RNA and peptide fragments within the cluster generally follows the same trend, except for the LYS mixtures, where RNA shows a higher tendency for aggregation (Figure 1C). Group P+ residues are considered the strongest promoters and form the largest cluster size and highest cluster density. Group P promoters still form relatively large clusters but are quantitatively less dense clusters compared to Group P+. The Group S suppressors form smaller, more loosely packed clusters. Finally, Group S+: strong suppressors form by far the smallest cluster sizes and lowest densities for the same biomolecular concentration of the mixture (Figure 1D). Notably, residues with longer side chains tend to suppress phase separation compared to their counterparts within the same side chain grouping (hydrophobic, polar, and charged). For instance, GLN exhibits lower phase separation affinity compared to ASN. Likewise, THR has lower phase separation affinity than SER, GLU is lower than ASP, and VAL is less than ALA. Interestingly, despite its long side chain, LEU shows a higher propensity to phase-separate primarily through hydrophobic interactions (Figure 2D). However, it does form the least dense clusters of Group S+ due to the steric impediments of its long side chain (Figure 1B-C).

Solvent accessible surface area (SASA) characterization within the cluster provides more insight into the size and density of phase-separated clusters. Considering the varying side chain lengths, SASA per atom offers a clearer depiction of RNA and residue packing within each phase-separated cluster. Using GLY as a reference, we note that hydrophobic and negatively charged residues tend to exhibit higher SASA per atom, while polar, aromatic, and positively charged residues demonstrate lower SASA per atom, indicating stronger interactions and denser clusters (Figure 1E). Residues ARG and TRP, despite their larger side chains, are found to possess the least SASA due to multivalent interactions leading them to be buried deep inside clusters.

To gain further insight into the microscopic driving forces of peptide-RNA phase separation, we have quantified atomistic contact frequencies of RNA-RNA, RNA-peptide, and peptide-peptide molecules inside clusters. The contact frequencies were then compared to the residues grouping defined by cluster size distributions (Figure 2A). Interestingly, this grouping perfectly matches the classification of residues according to their phase-separation propensity (Figure 1D). Group P+ (super promoters) form the highest RNA-GXG and GXG contacts, followed by Group P: Promoters. Groups S: Suppressor and S+: Super Suppressors have the lowest RNA-GXG and GXG contacts. We further dissected the three main types of interactions within systems: hydrogen bonds, pi-stacking, and hydrophobic interactions (Figure 2B-D & S6-S7).

Hydrogen bonds play a significant role in the LLPS of ARG, LYS, ASN, SER, GLN, and THR due to their higher number of -NH and -OH groups (Figure 2B). While HIS, TYR, and TRP facilitate phase separation via hydrogen bonds paired with pi-stacking and hydrophobic interactions, and PHE facilitates LLPS through pi-stacking and hydrophobic interactions alone (Figure 2B-D). Negatively charged residues (ASP and GLU) lead to suppression of LLPS. LEU exhibits phase separation through its hydrophobic interactions (Figure 2D).

We have computed the radial distribution function between functional groups of peptide and RNA fragments further to dissect preferential interactions between peptide and RNA fragments. We use the side chain and backbone of peptides as distinct functional groups.

Similarly, RNA is broken down into phosphate, sugar, and base functional groups. As expected, the positively charged residues like ARG and LYS predominantly interacted via their side chains with phosphate groups (Figure 3 and Figure S8A). On the other hand, polar residues ASN, GLN, SER, and THR exhibited interactions primarily with RNA bases, though they also showed significant interactions with phosphate and sugar groups (Figure 3 and Figure S8B-D). As expected, aromatic residues like PHE, TYR, TRP, and HIS clearly prefer pi-stacking interactions between their side chains with RNA bases rather than phosphate and sugar groups (Figure 3 and S9A-D).

**Figure 2.**
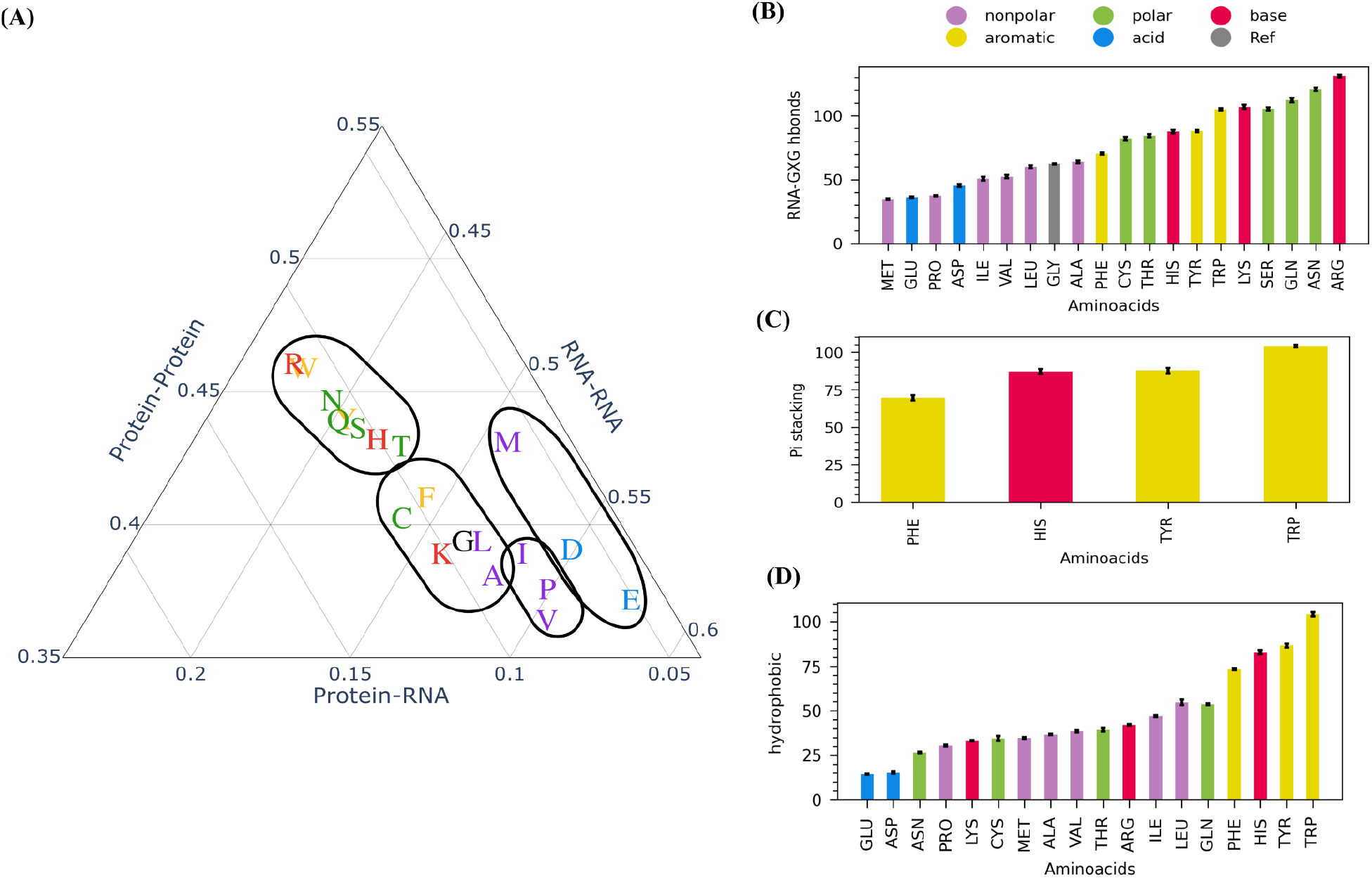
Atomistic profiles of interaction patterns of [U]_3_ and GXG peptide fragments. (A) A normalized ternary plot containing RNA-peptide, RNA-RNA, and peptide-peptide interaction probabilities for all 20 residues. (B) Number of hydrogen bonds between RNA and GXG for all 20 residues. (C) Number of pi-stacking interactions between RNA and GXG for all 20 residues. (D) Number of hydrophobic interactions between RNA and GXG for all 20 residues. Interaction values in B-D are shown as interactions per fragment.

**Figure 3.**
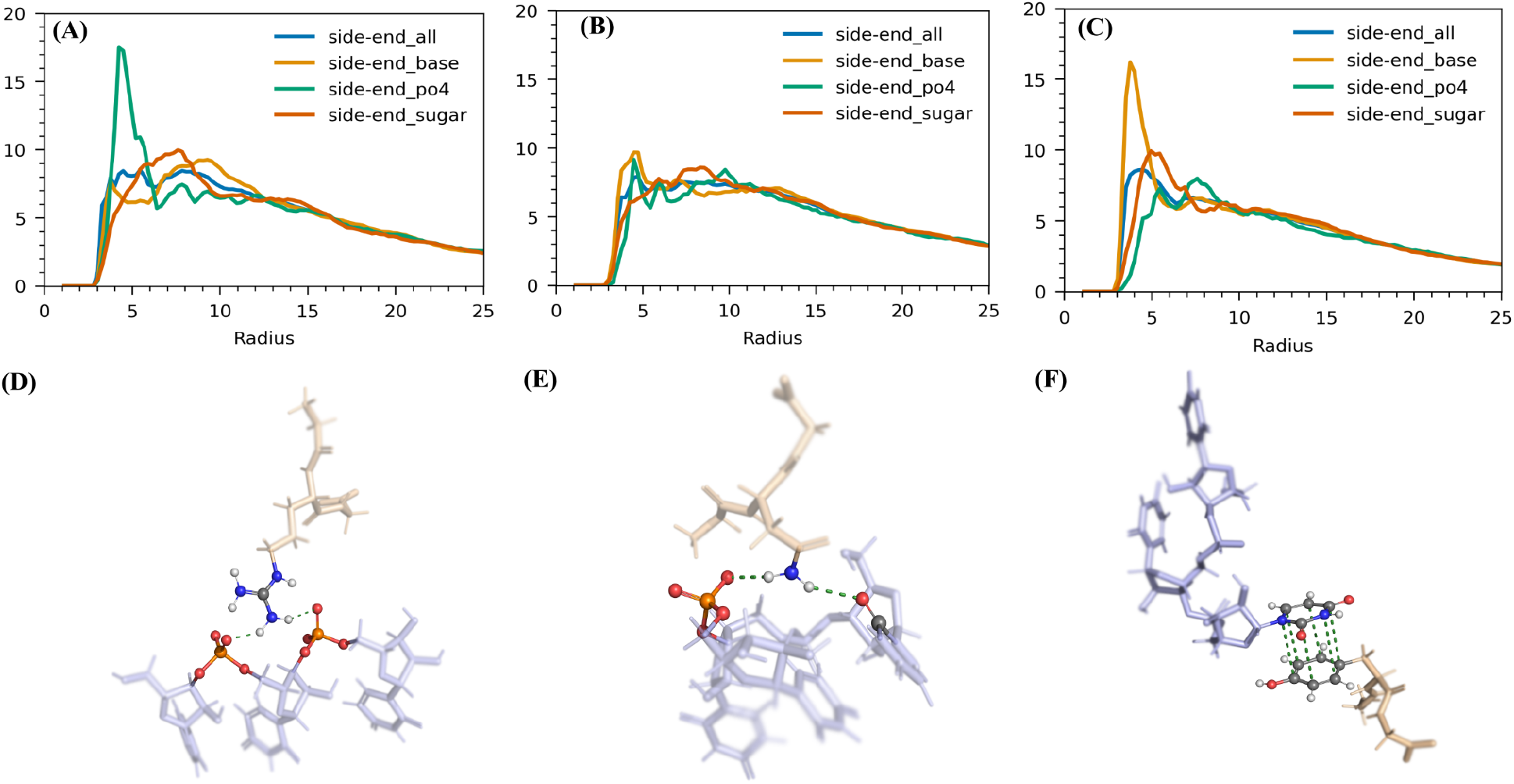
Specific interactions of RNA and peptide. Radial distribution function (RDF) of residue side chains with different parts of RNA (phosphate, sugar, and base) for various types of residues: (A) ARG showing a stronger preferential interaction with phosphate via electrostatic interactions, (B) ASN showing a balanced interaction with the base, sugar, and phosphate of RNA, and (C) TYR showing a stronger preferential interaction with the base via pi-stacking. The peptides are depicted in yellow, while [U]_3_ fragments are shown in blue, with their specific interactions highlighted.

Hydrophobic residues with larger side chains, such as ALA, VAL, ILE, LEU, and MET suppress cluster formation by disrupting some polar and hydrogen bond networks in clusters (Figure S2 and S10). Interestingly, LEU was found to interact with RNA bases through hydrophobic interactions (figure 2D). In the case of negatively charged residues, the RDF around the RNA is greater for the backbone than the side chain, suggesting that these residues’ side chains hinder interactions with RNA. ASP and GLU primarily interacted through the peptide’s backbone (Figure S11).

To assess the role of hydration in the phase separation of peptide and RNA fragments, we quantified water content around the cluster, focusing specifically on counting water molecules around RNA and peptide fragments within each cluster. We observed that water interacts more frequently with peptide fragments than RNA bases, which is consistent with our earlier work on the stoichiometric influence on RNA-peptide condensation [5]. As anticipated, residues prone to phase separation exhibit fewer water interactions, while those that suppress phase separation demonstrate higher water interaction levels (Figure 4A). Notably, in the system involving LYS, there is an unexpectedly low number of water molecules around RNA compared to LYS alone, elucidating why RNA tends to aggregate exclusively in LYS systems (Figure 4A). This trend is also observed in ALA systems. Water facilitates RNA-peptide phase separation by acting as a bridge between fragments. We quantified the number of water bridges where RNA and peptides are connected by either one (wb1) or two (wb2) water molecules. Charged residues such as ARG, LYS, ASP, and GLU exhibit a higher number of wb1 and wb2 compared to others (Figure 4B-E)

**Figure 4.**
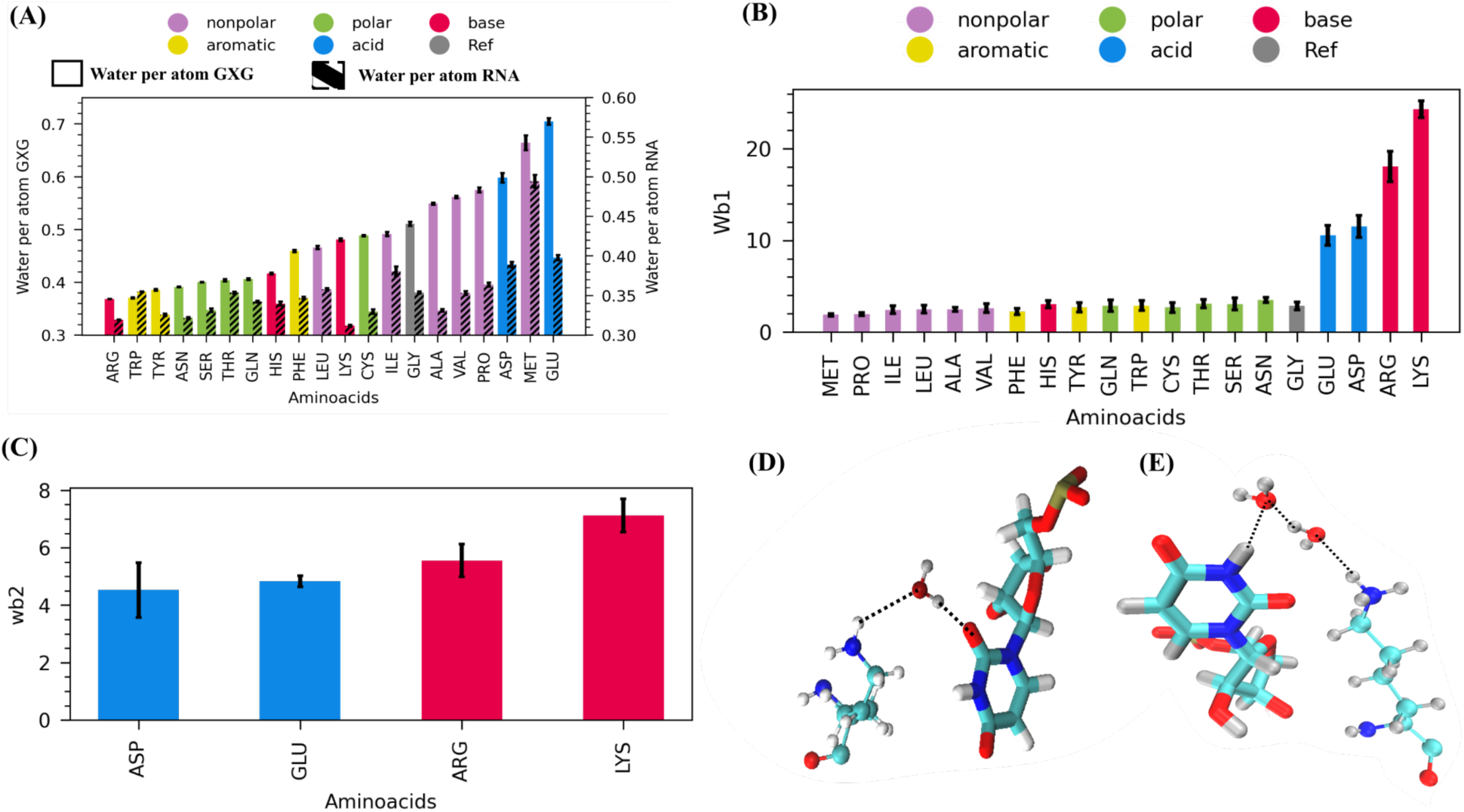
Water interactions with RNA and peptide clusters for all twenty residues. (A) The average number of water molecules per RNA (shaded region) and peptide fragment within the cluster. (B) The number of single water bridges (WB1) connecting RNA and GXG. (C) Number of double water bridges (WB2) connecting RNA and GXG. The schematic representation of (D) WB1 and (E) WB2 illustrates single and double water bridge formations between RNA and peptide fragments.

Finally, we examined the interaction of ions, particularly Na+, with RNA in different phase-separated mixtures. We observed that RNA drives Na+ ions out of clusters most strongly in mixtures containing LYS and ARG due to their interaction with the phosphates (Figure S3 A-B). Interestingly, in residues such as ASN and GLN, there is a notable increase in Na+ ion interaction with RNA, indicating that ions can also play a significant bridging role to help promote cluster formation and growth (Figure S3 B). Additionally, Cl-ions were found to be abundant around RNA in LYS and ARG systems due to their positively charged side chains.

ASN and GLN tend to repel Cl-more than charged residues ASP and GLU (Figure S4 B). To conclude our study has elucidated the chemically specific contacts, hydration profiles, and ionic effects that can either drive or inhibit protein-RNA phase separation. We have elucidated the chemically specific contacts, hydration profiles, and ionic effects, illuminating the driving forces that promote or suppress protein-RNA phase separation. We find that our results are best rationalized by creation of four categories for protein residues based on their ability to promote, maintain, suppress, or disrupt protein-RNA clusters. By mapping a ternary phase diagram, we demonstrate that the promotion or suppression of liquid phases results from the intricate interplay of RNA-RNA, peptide-RNA, and peptide-peptide interactions. Future studies will be needed to probe deeper into the role of context-specific sequence effects and the role of RNA sequence by examining more complex sequence patterns that still need to be addressed in the present study.

## Computational Methods

Twenty individual simulations were run, each containing 1 type of GXG peptides (uncapped terminal glycines with X being one of the 20 standard amino acids) and [U]_3_ RNA fragments. The PDB structures of the peptides were produced using PeptideBuilder [30], and that of the [U]_3_ RNA fragments was obtained from X3DNA [31,32]. These biomolecules were then packed into the system at a 1:1 mass ratio using packmol [33], creating a biomolecular system density of ∼100 mg/mL inside a ∼100 Å cubic box, and solvated using explicit TIP3P water model [34] paired with the amber99SB*-ILDN-Q forcefield [35,36]. Previous computational work[28,29] on short peptide mixtures have shown that TIP3P water model is optimal choice for water to ensure phase separation. A NaCl concentration of 100 mM was used for each simulation. Once each system was packed and solvated, they were run through an energy minimization, a 100ns NVT ensemble, and equilibration, followed by a 2 μs productive run in the NPT ensemble using OpenMM’s Monte Carlo barostat and Langevin Middle integrator to maintain a constant temperature of 300K and a constant pressure of 1 atm. A 2 fs time step was used throughout each of the simulations.

The convergence of each of the systems was confirmed via RDF functions as a function of time between peptide and RNA fragments (Figure S1). The DBSCAN algorithm was used to extract the clusters in a frame-by-frame analysis to analyze the size, density, and clustered residue fractions of the clusters formed from the different peptides with the RNA fragments. A detailed contact analysis was also performed using Prolif [37] to investigate the linkages between the cluster characteristics and the various interactions (hydrophobic, electrostatic, and water-bridged interactions) responsible for cluster formation and maintenance. Water-biomolecule interactions were quantified by extracting the number of water molecules in direct contact with both peptide and RNA fragments inside the clusters. A detailed water-bridged interaction analysis was also performed using MDAnalysis [38,39]. Lastly, a quantitative analysis of Na+ and Cl-ions was performed by simply tracking the number of each ion inside the cluster throughout the simulations.

## ACKNOWLEDGMENTS

Authors acknowledge support from the National Institutes of Health with grant no R35GM138243.

## For Table of Contents Only

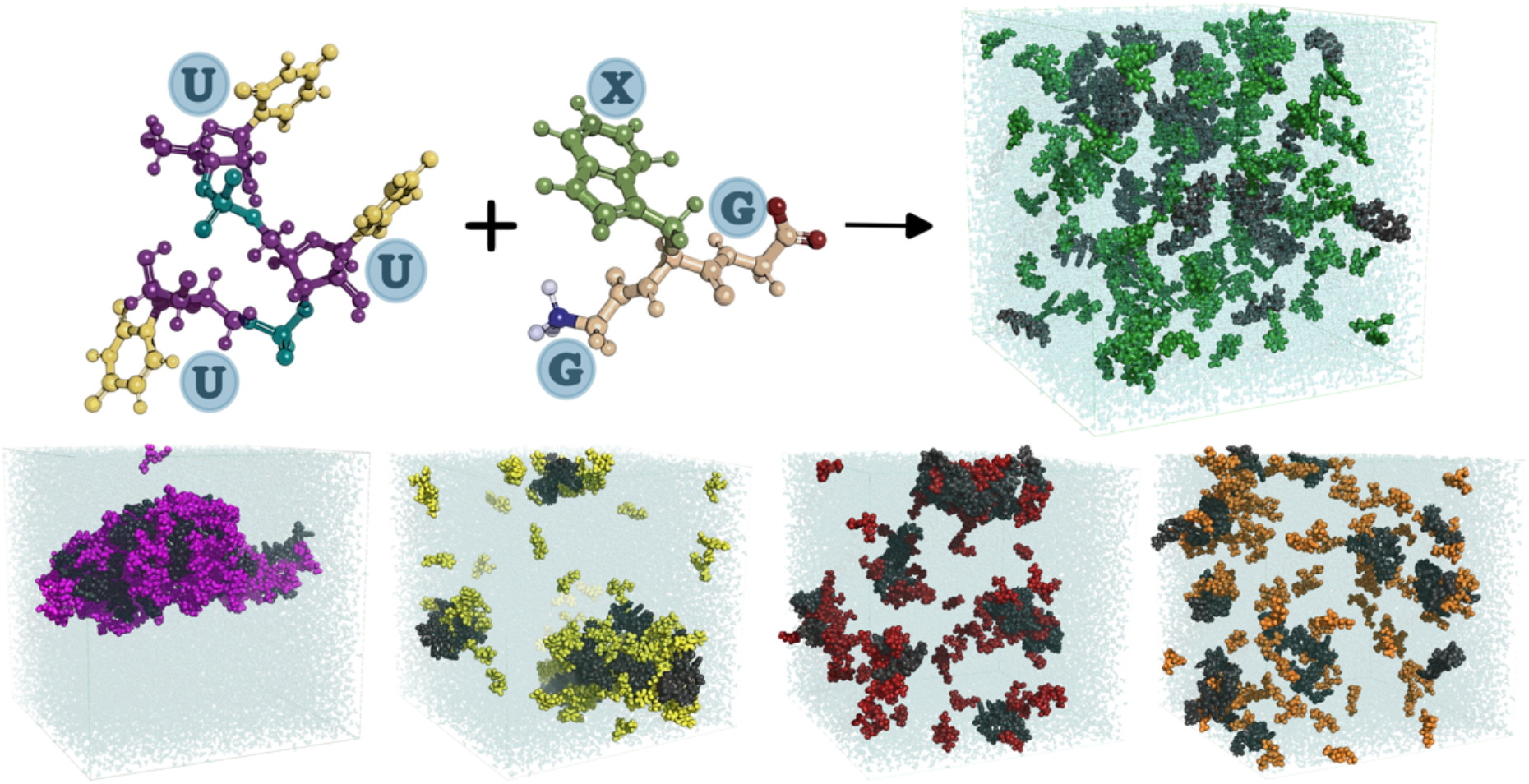

## Notes

### Competing Interest Statement

The authors have declared no competing interest.

