## Supplemental Information for "Nucleoprotein phase-separation affinities revealed via atomistic simulations of short peptide and RNA fragments"

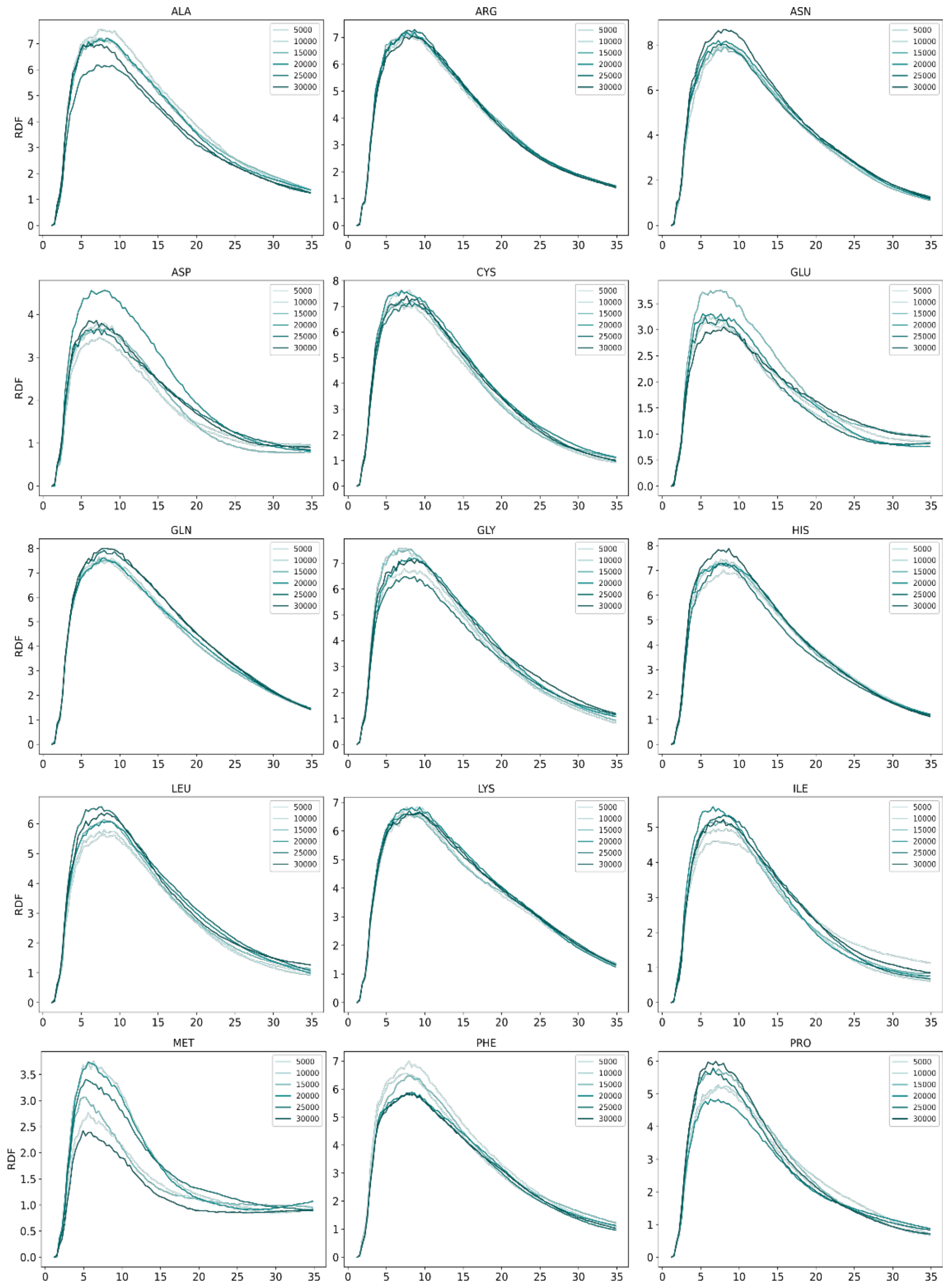


**Figure S1:** Radial distribution function of U3 and GXG for all twenty amino acids as a function of time, showing the convergence of the simulation over time.

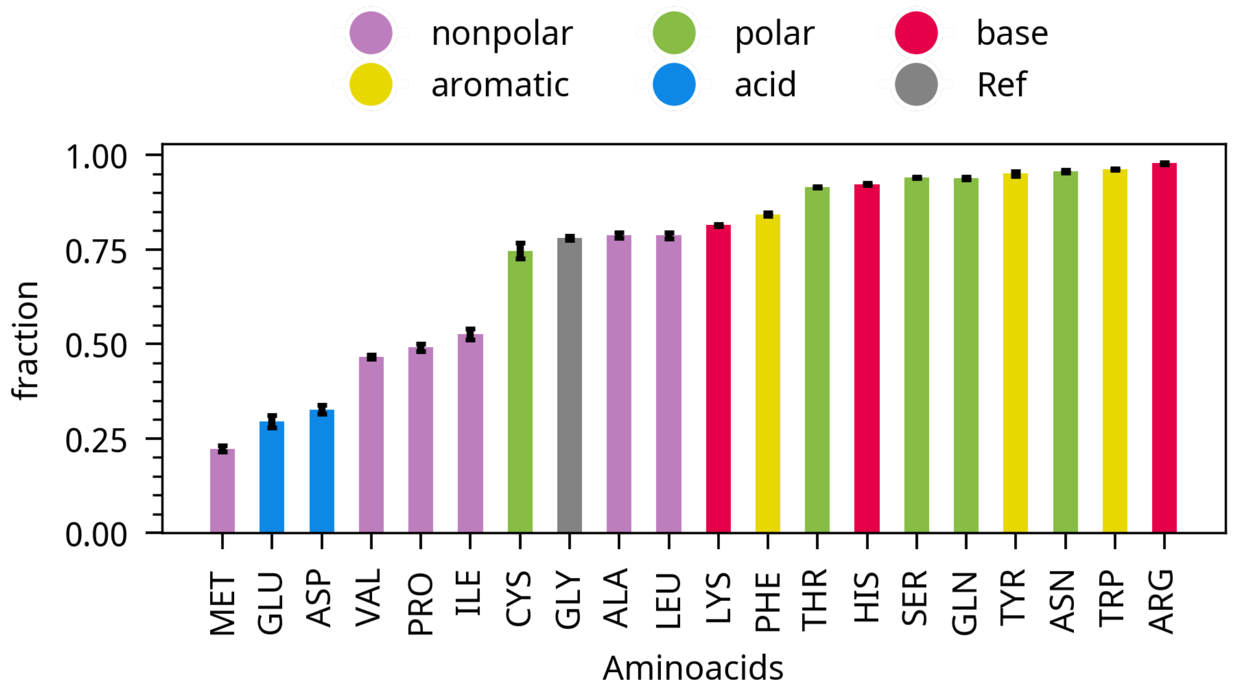


**Figure S2:** Cluster size quantified by fraction of biomolecules (U3 and GXG)

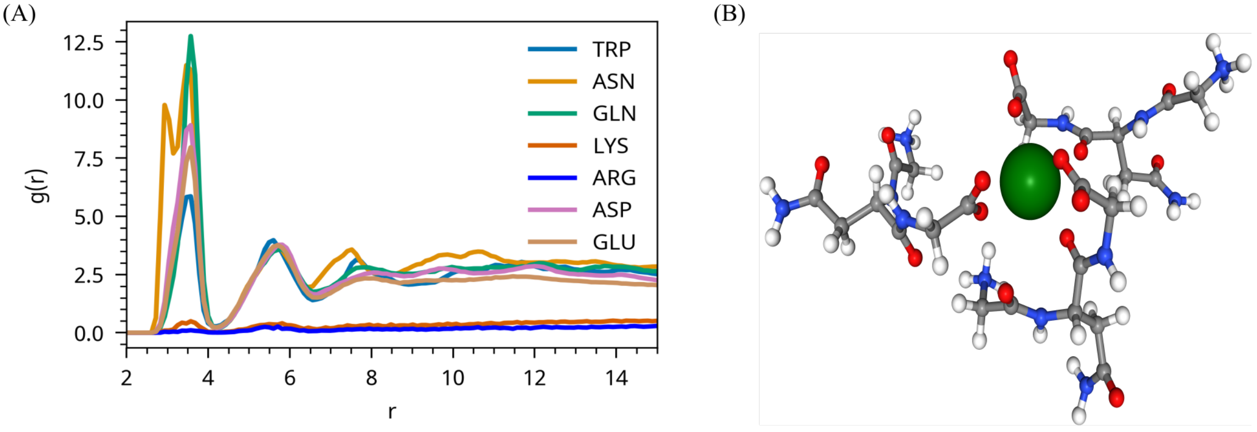


**Figure S3:** Interaction of ions with RNA, quantified using radial distribution functions (RDF). (A) Interaction of Na+ ions with RNA, showing the distribution and affinity of sodium ions around the RNA molecule. (B) Snapshot of Na+ interacting with ASN.

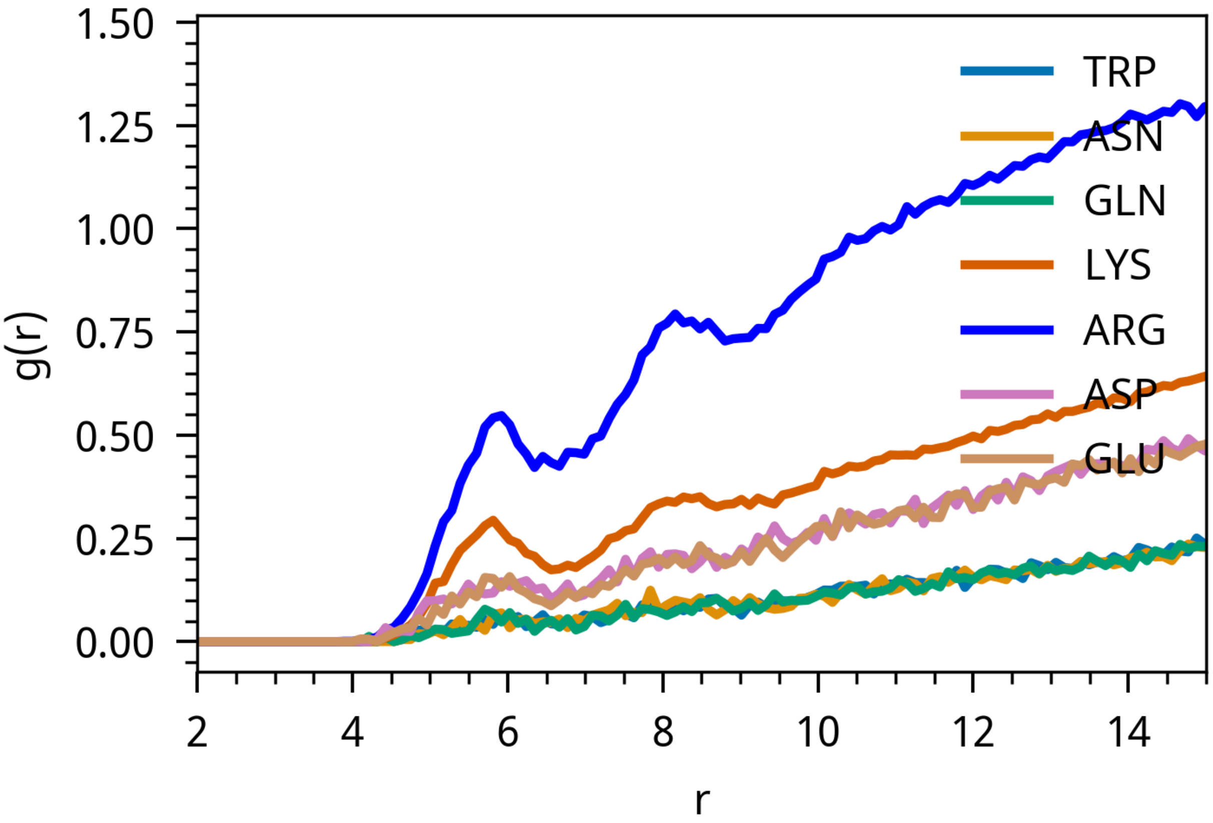


**Figure S4:** Interaction of Cl- ions with RNA, illustrating the distribution and affinity of chloride ions around the RNA molecule.

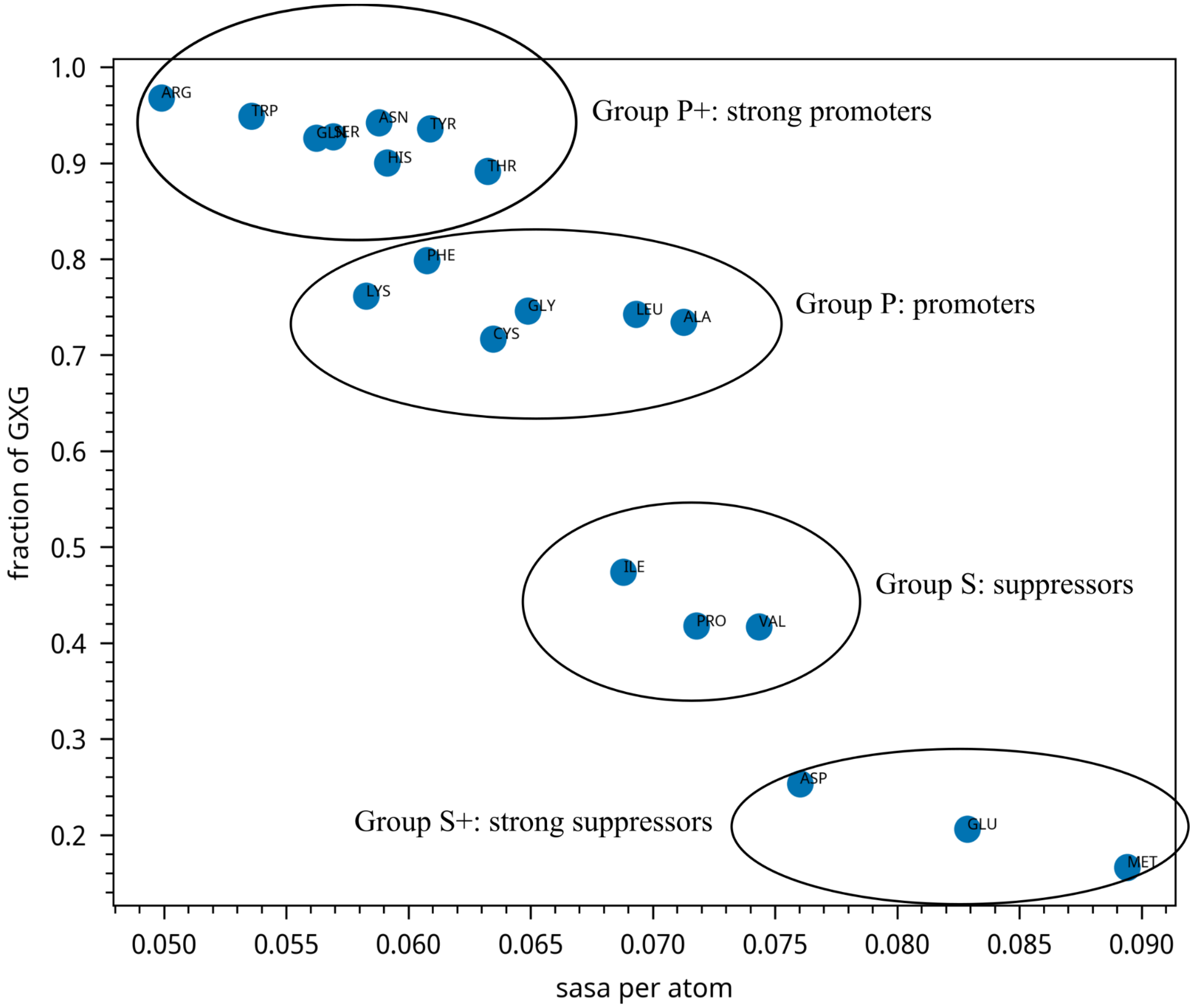


**Figure S5:** Solvent accessible surface area (SASA) per atom vs. cluster size for all 20 residues calculated from 20 simulations of GXG-RNA mixtures.

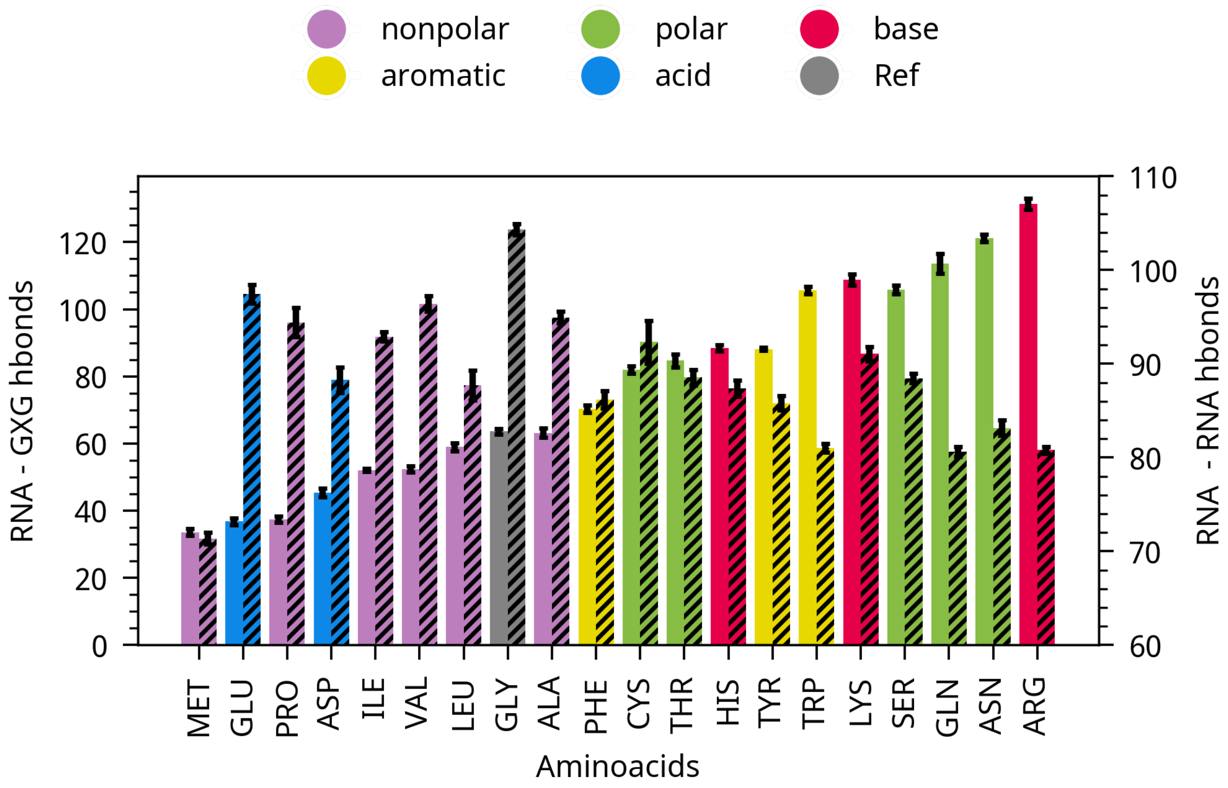


**Figure S6:** RNA-protein hydrogen bonds (shown in normal bar) and RNA-RNA hydrogen bonds (shown in shade bar).

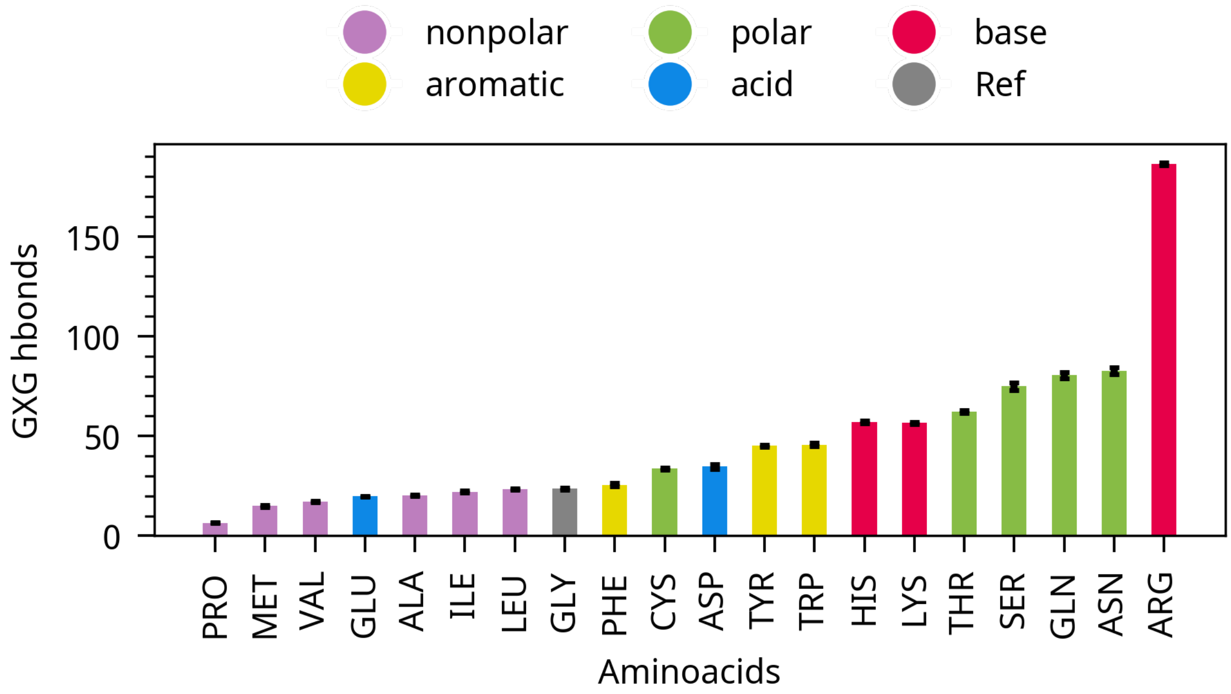


**Figure S7:** Number of hydrogen bonds between GXG peptides

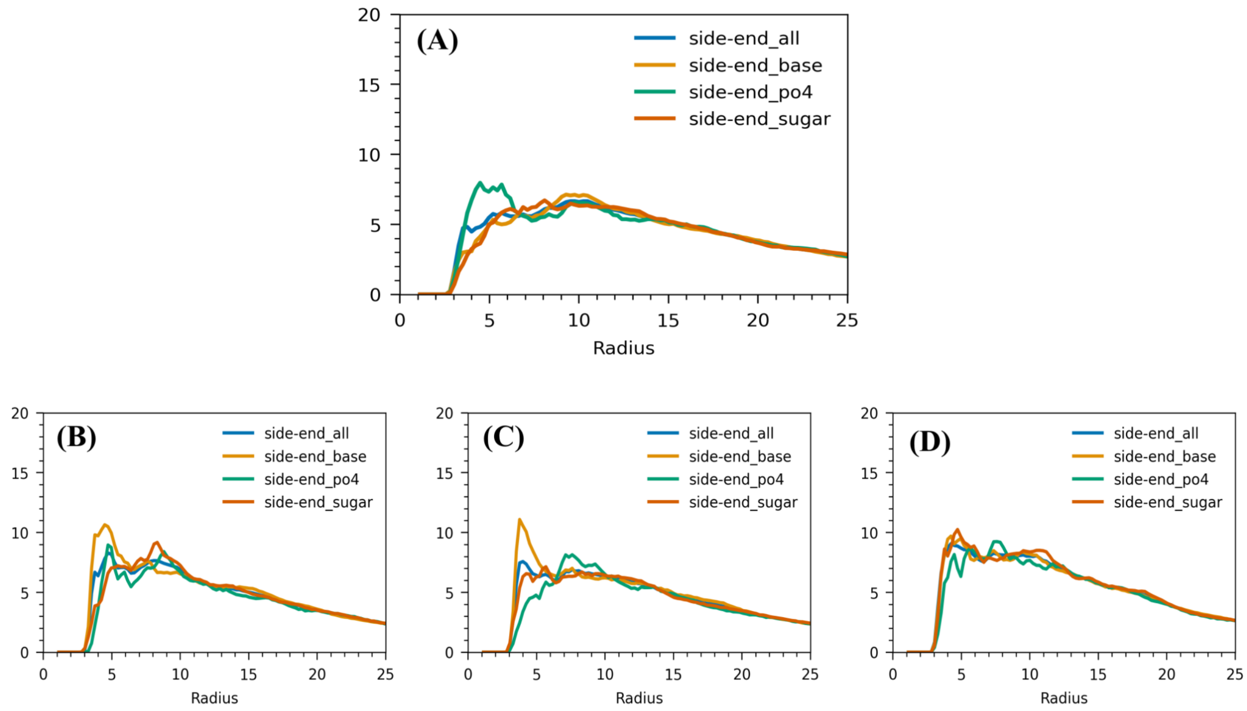


**Figure S8:** Radial distribution function (RDF) of residue side chains with different parts of RNA (phosphate, sugar, and base) for various types of residues: (A) LYS , (B) GLN (C) THR and (D) SER

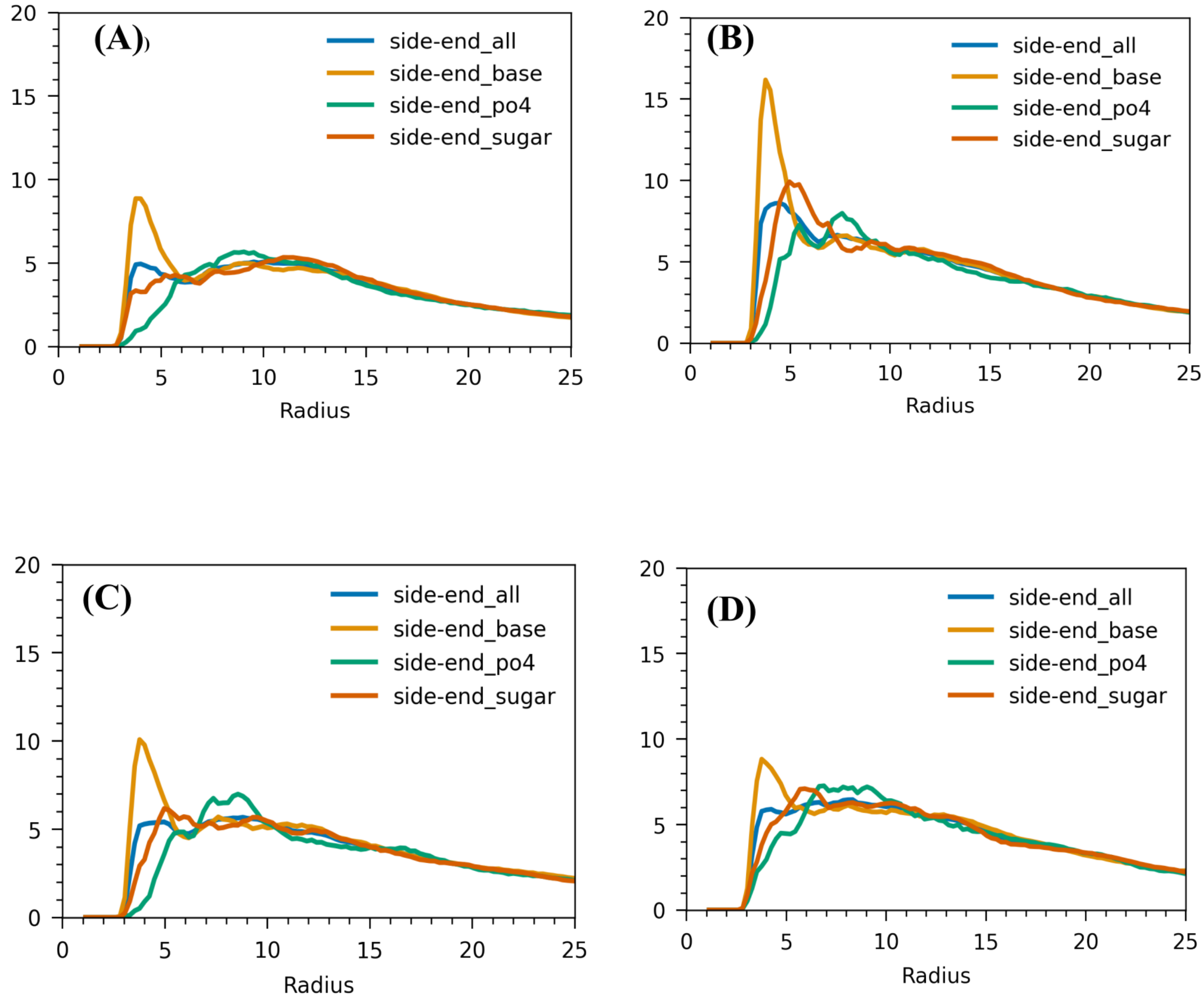


**Figure S9:** Radial distribution function (RDF) of residue side chains with different parts of RNA (phosphate, sugar, and base) for various types of residues: (A) PHE, (B) TYR (C) TRP and (D) HIS

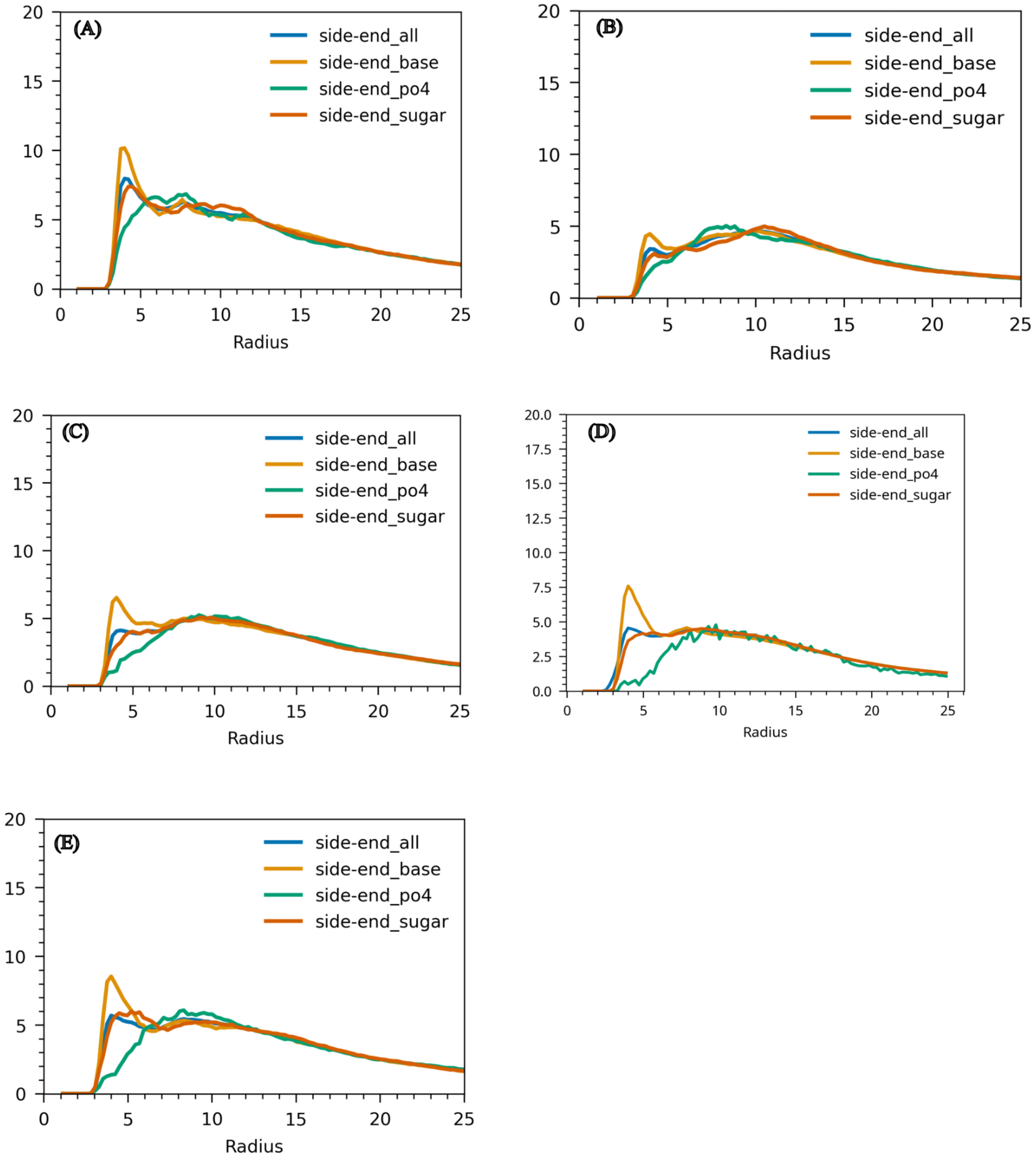


**Figure S10:** Radial distribution function (RDF) of residue side chains with different parts of RNA (phosphate, sugar, and base) for various types of residues: (A) ALA , (B) VAL, (C) LEU (D) ILE and (E) MET

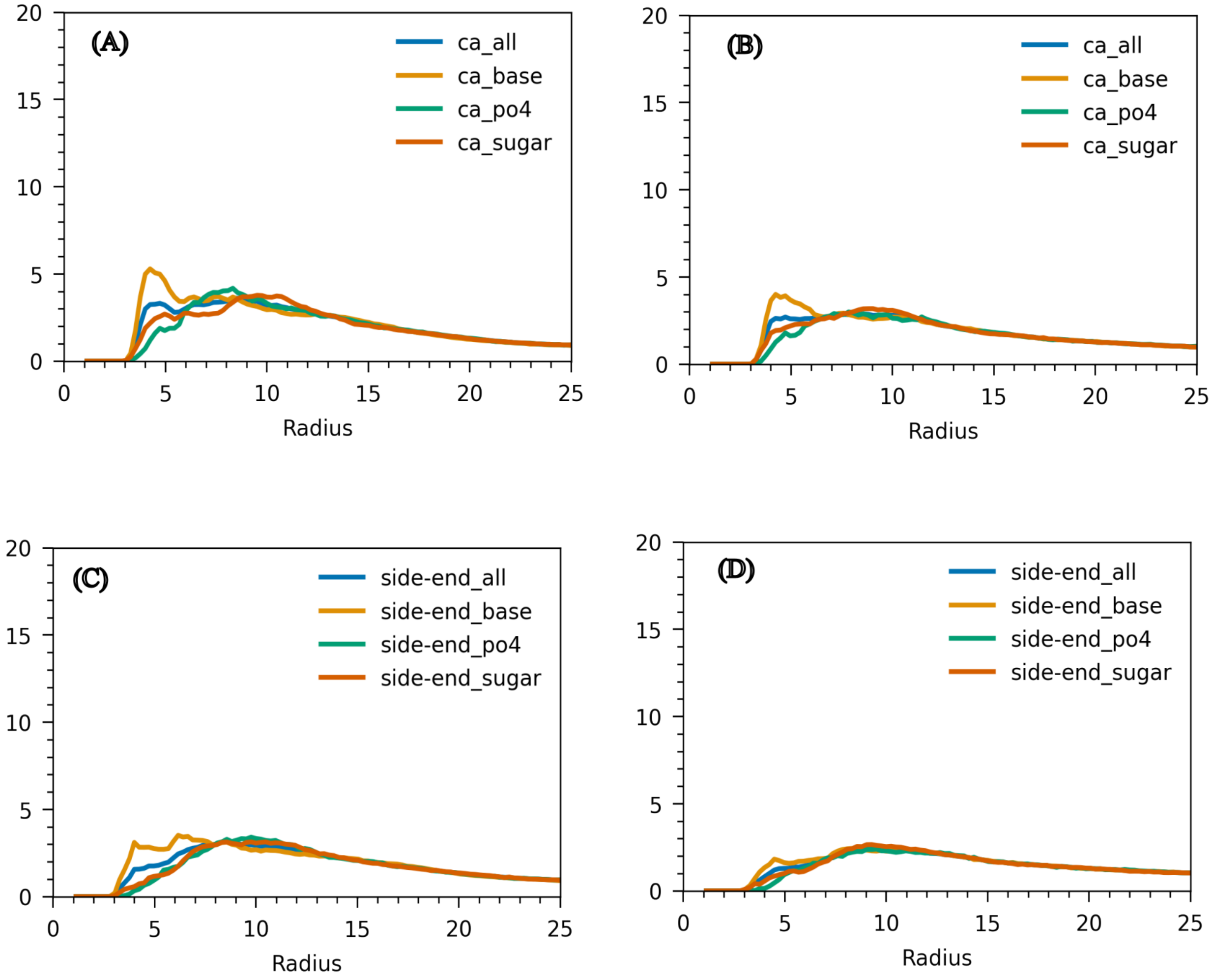


**Figure S11:** Radial distribution function (RDF) of residue with different parts of RNA (phosphate, sugar, and base) for various residues; the residue is divided into CA and side chain end. CA and different parts of RNA of (A) ASP and (B) GLU. side end of residue and different parts of RNA of (C) ASP and (D) GLU. Overall it shows side end suppress phase separation of residues.
